# Investigating on-road lane maintenance and speed regulation in post-stroke driving: a pilot case-control study

**DOI:** 10.1101/549410

**Authors:** Heng Zhou, Qian (Chayn) Sun, Alison Blane, Brett Hughes, Torbjörn Falkmer, Jianhong (Cecilia) Xia

## Abstract

The post-stroke population is expected to rise and the cohort of post-stroke individuals who wish to return to driving is likely to rapidly increase too. Stroke can adversely affect coordination and judgement of drivers due to the executive dysfunction, which is relatively common in the post-stroke population but often undetected. Quantitatively examining vehicle control performance in post-stroke driving becomes essential to inspect whether and where post-stroke drivers are risky. To date, it is unclear as to which indicators, such as lane keeping or speed control, can differentiate driving performance of post-stroke drivers from that of normal (neurotypical) older drivers. By employing a case-control design using advanced vehicle movement tracking and analysis technology, this pilot study aimed to compare the variations in driving trajectory, lane keeping and speed control between the two groups using spatial and statistical techniques. The results showed that the mean Standard Deviation of Lane Deviation (SDLD) in post-stroke participants was higher than those of normal participants in complex driving tasks (U-turn and left turn) but almost the same in simple driving tasks (straight-line sections). No statistically significant differences were found in the speed maintenance between the post-stroke drivers and the normal adults. The findings indicate that, although older drivers can still drive as they need to after a stroke, the decline in cognitive abilities still imposes higher cognitive workload and more effort for post-stroke older drivers. Future studies can investigate post-stroke adults’ driving behaviour at more challenging driving scenarios, or design driving intervention program to improve their executive function in driving.

## 1. INTRODUCTION

Driving is an important, autonomous daily activity, which underpins personal mobility in society (Burns, 1999). It is a vital skill that requires the use of multiple neuropsychological processes such as cognitive, visual, perceptual and data processing abilities (Anstey, Wood, Lord, & Walker, 2005).

It has been well-reported that older adults tend to experience cognitive decline with increased age (Mathias & Lucas, 2009; Q. C. Sun, Xia, Foster, Falkmer, & Lee, 2018b) and this decline can be accentuated by the neural brain damage caused by a stroke (Motta, Lee, & Falkmer, 2014; Poole, Chaudry, & Jay, 2008), thereby subsequent adverse impact on their driving ability (Harbluk, Noy, & Eizenman, 2002). This is a cause for concern given the risk of a stroke increases with age and that the proportion of older adults continuing to drive is increasing (Howard & Goff, 2012). Therefore, as the post-stroke population is expected to rise to over 132,000 by 2050 in Australia (National stroke foundation, 2014), the population of post-stroke individuals who wish to return to driving is likely to rapidly increase (Terblanche & Wilson, 2015).

In consequence, ensuring a safe return to driving through accurate driving measurements is required, particularly as the older driver population, and particularly post-stroke drivers, are at an increased risk of a crash and of fatal injuries due to increased frailty (McGwin, Sims, Pulley, & Roseman, 2000; M. Rabadi, A. Akinwuntan, & P. Gorelick, 2010; Rakotonirainy, Steinhardt, Delhomme, Darvell, & Schramm, 2012).

Currently, driving simulators and on-road driving assessments are the two main methods used to assess driving behaviours, such as lane keeping performance (Cao & Liu, 2013a; He, McCarley, & Kramer, 2014). For the simulated driving scenarios, the participants perform a lane maintaining task and their lane deviation is calculated automatically by a driving simulator program (He et al., 2014). In contrast, on-road assessments involve completing a variety of driving tasks on-road, in a licensed vehicle and these are considered the ‘gold standard’ because they take place in the real world, and are therefore more likely to gain a more accurate representation of drivers’ performance (Carsten, Kircher, & Jamson, 2013; Selander, Lee, Johansson, & Falkmer, 2011). Although preferable as an assessment method, it is hard to detect subtle variations between drivers in an on-road assessment if they do not make any errors as traditional driving assessments rely on subjective observations from driving evaluators. Anstey et al. (2005) pointed out that using a fixed driving route for on-road testing is an efficient strategy to examine the participant’s driving performance. Therefore, the optimal approach is to record their driving in naturalistic settings at a microscopic level using a fixed driving route. Advanced Global Navigation Satellite System, such as GPS technology, can be applied to tracking vehicle movements and trajectory, which can be used to ascertain the driver’s driving behaviour on road assessment.

The driving trajectory and behaviours are both dependent on the position data from the Global Positioning System (GPS), thus the accuracy and precision of the GPS data becomes the key point in analysing the driver’s driving trajectory and behaviours. Recently, multiple satellite systems have contributed significantly to global navigation and positioning systems regard to precision and availability. Generally, the combination of Multi-Global Navigation Satellite System (GNSS) receivers can be used to collect and detect a driver’s driving trajectory (Q. Sun, Xia, Foster, Falkmer, & Lee, 2016; Q. C. Sun, Xia, Foster, Falkmer, & Lee, 2018a). It can record the position data of a vehicle to one tenth of a second with a high degree of precision (1 decimetre), which is important for improving driving trajectory detection accuracy. In comparison with single GPS technology, Multi-GNSS can provide a better approach to recording more accurate position information since more satellites can be tracked, which will be efficient in tracking the drivers’ vehicle control movement trajectory. To improve the raw position data, corrections can be applied to the recorded position data when the accuracy is particularly important (Q. C. Sun et al., 2017). Specifically, efficient GPS position techniques can be used to correct errors in the position data. For example, Real time kinematic (RTK) can be applied to enhance the accuracy and precision of the position data which can be fitted to track driving trajectories. The RTK technique introduces not only GNSS code pseudo-ranges measurements to compute its position, and atmospheric errors such as troposphere errors and ionosphere errors could be considered and evaluated (El-Rabbany, 2006), it also applies carrier phase measurements, which can provide positions that are orders of magnitude. Therefore, the position data could reach an accuracy level of between a millimetre to a centimetre (Q. C. Sun et al., 2017).

By adopting the millimetre to a centimetre measure of driving trajectory using multi-GNSS RTK technologies, lane keeping performance can be assessed at a high accuracy level in a GIS (Geographic Information System) platform (Q. C. Sun et al., 2017).

Given the increasing population of post-stroke drivers and this new on-road technology, the primary aim of this study was to explore the quantitative variations in vehicle control performance between post-stroke and neurotypical (‘normal’ from now on) older drivers using advanced vehicle tracking and spatial analysis technologies. Based on the literature review above (Cao & Liu, 2013b; Perrier, Korner-Bitensky, Petzold, & Mayo, 2010; M. H. Rabadi, A. Akinwuntan, & P. Gorelick, 2010), the objectives of this paper are to examine the vehicle control performance in post-stroke driving in comparison with normal drivers’ performance through a pilot case control study, and to explore potential indicators of post stroke driving that can be employed in driving intervention design for this cohort drivers.

## 2. METHODS

### 2.1 Design

This pilot study employed a case-control design to measure and compare the differences in on-road lane maintenance, trajectory and speed control in post-stroke drivers and age-matched controls using advanced vehicle movement tracking and GIS technologies.

### 2.2 Participants

Participants comprised 14 post-stroke adults (79% male, M=71.1 years, SD=6.6) and 14 older drivers who had not had a stroke (57% male, M=72.9 years, SD=6.5) to act as a comparison group. The inclusion criteria for the study participation were that participants held a driving licence valid within Australia, had at least one year of overall driving experience, drove at least twice a week and had access to a fully insured vehicle. Further criteria for the post-stroke cohort were that they had previously been diagnosed with a stroke (either ischemic, haemorrhagic or a transient ischemic attack) and had been cleared to drive by a medical professional. Participants were excluded if they had been diagnosed with hemianopia, a neurodegenerative disease, such as Parkinson’s disease or dementia, and if they required a wheelchair to get around. Participants were recruited using volunteer sampling techniques such as using local community groups, post-stroke support groups, community newspapers and local radio stations.

### 2.3 Driving Measures

#### 2.3.1 Lane Keeping

Lane keeping performance is a vital indicator of driving performance (Cao & Liu, 2013b) and it is commonly measured by the Standard Deviation of Lane Deviation (SDLD), with lower standard deviation of lane deviation values indicating better lane keeping performance (Cao & Liu, 2013a). Lane deviation is defined as the perpendicular distance between the vehicle position and the lane centre line (Peng et al., 2013). Cao and Liu (2013a) suggested that lower driving speed can reduce the driver’s mean value of standard deviation of lane deviation (SDLD). Their statistical analysis found that the mean value reduced from 0.36 m to 0.30 m when the driving speed reduced from 72 km/h to 36 km/h. In addition, more difficult road geometry requires more complex mental cognition workload and executive function, which may lead to poor lane keeping performance. However, some studies also show that if the road geometry of a lane keeping task is quite simple the required mental workload becomes very low, which could lead to a poor lane keeping performance due to a lack of excitement and motivation (White, 1959). Therefore, this work focused on some of the more demanding parts of the on-road test’s driving route such as the roundabouts and left turns. Participants’ driving trajectories in simple driving tasks (straight driving) were also explored in this study.

#### 2.3.2 Speed Control

The capability of speed control can also influence the drivers’ on-road performance and driving safety. It is known that older and novice drivers are more likely to respond inappropriately to road geometries and traffic signs, which require them to control driving speed (Chan, Pradhan, Pollatsek, Knodler, & Fisher, 2010). Similarly, Weihong, Blythe, Edwards, Pavkova, and Brennan (2015) found that 80% of 60+ drivers did not control their speed appropriately in free-flow traffic conditions and older drivers have been found to commit more driving errors when compared to younger drivers, however their speed adjustment and lane position was improved in automatic transmission cars (Selander, Falkmer, & Bolin, 2012). Furthermore, older drivers usually experience high crash risk comparing to other cohorts in urban environments, due to their inappropriate speed control performance (Weihong et al., 2015). Hence, the mean speed and standard deviation of speed can indicate the drivers’ speed control performance (Reed & Green, 1999).

### 2.4 Driving data collection and processing

Data collection for this study took place during the normal traffic time. Participants were required to complete an on-road driving task with their own vehicle. A Trimble R10 Multi-GNSS receiver was attached to the roof of the vehicle recording moment-to-moment GPS data at 10 Hz, which records the vehicle positions at every 10 seconds.

To improve the raw position data, corrections were applied to the recorded position data. Following Q. Sun, Xia, Nadarajah, et al. (2016) efficient GPS position techniques were used to correct errors in the position data. The Real time kinematic technique (RTK) was applied to enhance the accuracy and precision of the position data, which was fitted to track driving trajectories. By adopting the millimetre to a centimetre measure of driving trajectory using multi-GNSS RTK technologies, lane keeping performance was assessed at a high accuracy level, therefore, providing a solid foundation for this comparative study of driving performance in normal and post-stroke older drivers (Q. C. Sun et al., 2018a).

### 2.5 Procedure

The on-road driving task was completed within and around Curtin University, Bentley, in Western Australia. The on-road assessment route (Appendix A) was specifically selected to contain intersections, roundabouts, U-turns, traffic lights. Participants were unaware of the route prior to the assessment and were given directions as the route progressed, for example, the drivers were required to notice and react to different speed limits. The whole route was approximately 9 km long containing various speed limits and traffic scenarios and took approximately 20 minutes to complete. Data were collected for the duration of the assessment route and also data from specified points during the route were then processed and mapped using ArcGIS software, following the procedure of Q. Sun, Xia, Nadarajah, et al. (2016). The residual output provided accurate vehicle coordinates used to calculate SDLD and speed control across all participants in both groups.

### 2.6 Data Analysis

The foci of this study were the lane maintenance; speed control and driving trajectory measured whilst undertaking a U-turn at a roundabout, a left turn and two straight line sections (as indicated in Appendix A). The driving trajectory of each participant consisted of a set of position points at every 10 seconds inteval. The perpendicular distances between the points and the benchmark line were defined as the lane deviations. The perpendicular tool in ArcGIS software was applied to draw and calculate the distance of the perpendicular segments between the vehicle’s position points and the benchmark line (Q. C. Sun et al., 2018a).

The SDLD and speed in each driving task (e.g., U-turn, left turn and straight-line section) were also calculated. Figure 2 illustrates one example of lane deviation (distance of the perpendicular segments) in a U-turn based on the perpendicular tool. Due to the geometry and complexity of the U-shaped roundabout, the roundabout was divided into three parts, entering U-turn/entry, middle part and exiting U-turn/exit (see Figure 2). Therefore, the lane keeping performance was analysed in each part of the roundabout.

**Figure 1:**
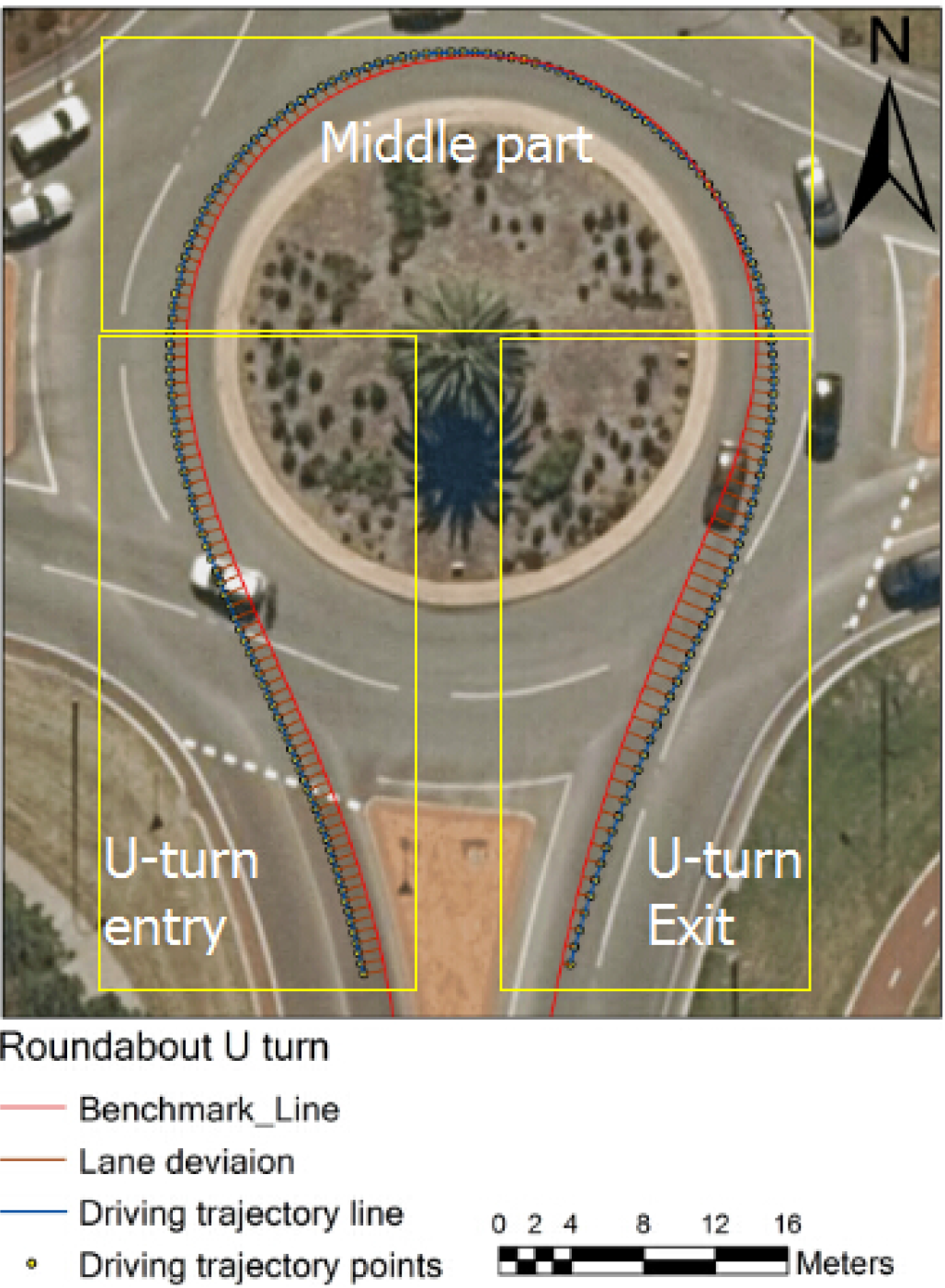
Driving trajectory and lane deviation from benchmark line in a U-turn and the three sections of the U-turn

Data were tested for normality with the Kolmogorov-Smirnov test and were found to be normally distributed. One way ANOVA test in SPSS version 21.0 (IBM Corporation, 2012) software was applied to examine differences of SDLD between the post-stroke group and normal comparison group in both complex (driving through a U-turn and left turn) and simple driving tasks (straight line driving). The critical *α*-value was set to 0.05 for all tests. With 28 participants providing data, a Cohen’s *d* of 1.03 was possible to detect at a 1-*β* of 0.8.

### 2.7 Ethical considerations

This research and the associated study protocols were approved by the Curtin University Human Research Ethics Committee (approval number: HR206-2014 and HR68-2014).

Participants were presented with an information sheet, given the opportunity to ask questions. Each participant provided signed informed consent prior to participation. Participants were also informed that they could leave the study at any time without incurring any negative consequences.

## 3. RESULTS

The results were generated based on spatial and statistical analyses of the differences of lane keeping performance between the post-stroke and neurotypical (normal) older drivers.

### 3.1 Spatial analyses of driving trajectories across the groups

Figure 3 shows the post-stroke and normal drivers’ vehicle movement trajectories in the exit part of a U-turn. The post-stroke older drivers’ deviation from the road centre line in U-turn exit was larger than that of normal older drivers. The recorded driving trajectory of the post-stroke older drivers did not cluster around the centre line of the road in U-turn exit; furthermore, some participants merged to another lane during driving through the exit part of the U-turn. Comparatively, all the normal older drivers’ driving trajectories distributed more closely towards the benchmark line, as demonstrated in Figure 3, suggesting that the post-stroke drivers’ lane keeping performance in a relatively complex driving task was deemed poorer than that of normal drivers.

**Figure 2:**
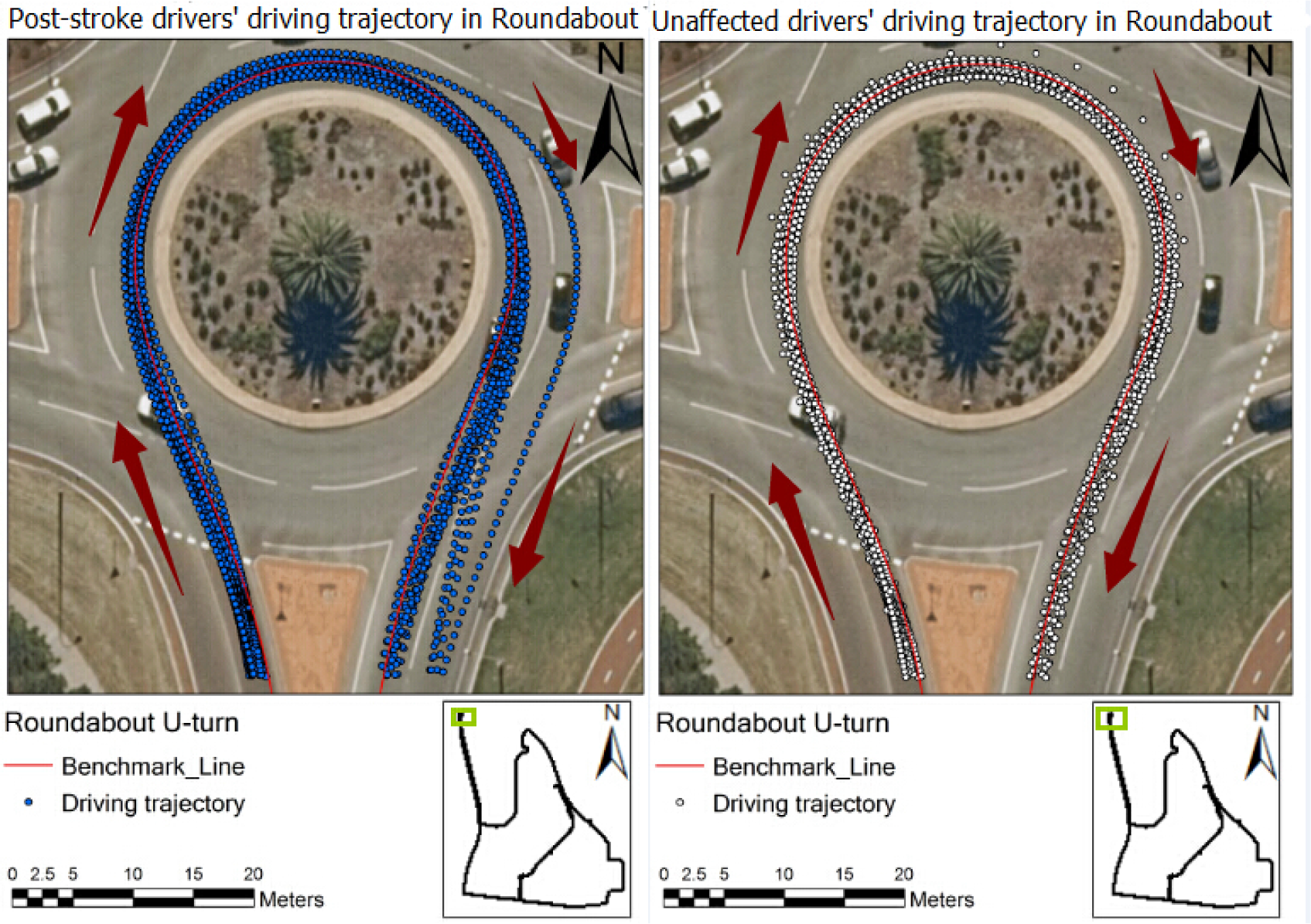
Driving trajectory of post-stroke (left) and normal (right) groups in a U-turn.

Figure 4 exhibits the post-stroke drivers’ and normal drivers’ vehicle control trajectories in the straight-line section of the on-road test. The blue points in Figure 4 represent the post-stroke older drivers’ vehicle control trajectories, which were gathered on the centre line of the lane. Surprisingly, the distances (lane deviation) between the normal drivers’ vehicle control trajectory and the benchmark line in Figure 4 were relatively larger than those of post-stroke drivers, indicating that the normal older drivers’ lane keeping performance in simple driving tasks may be actually poorer than that of the post-stoke drivers.

**Figure 3:**
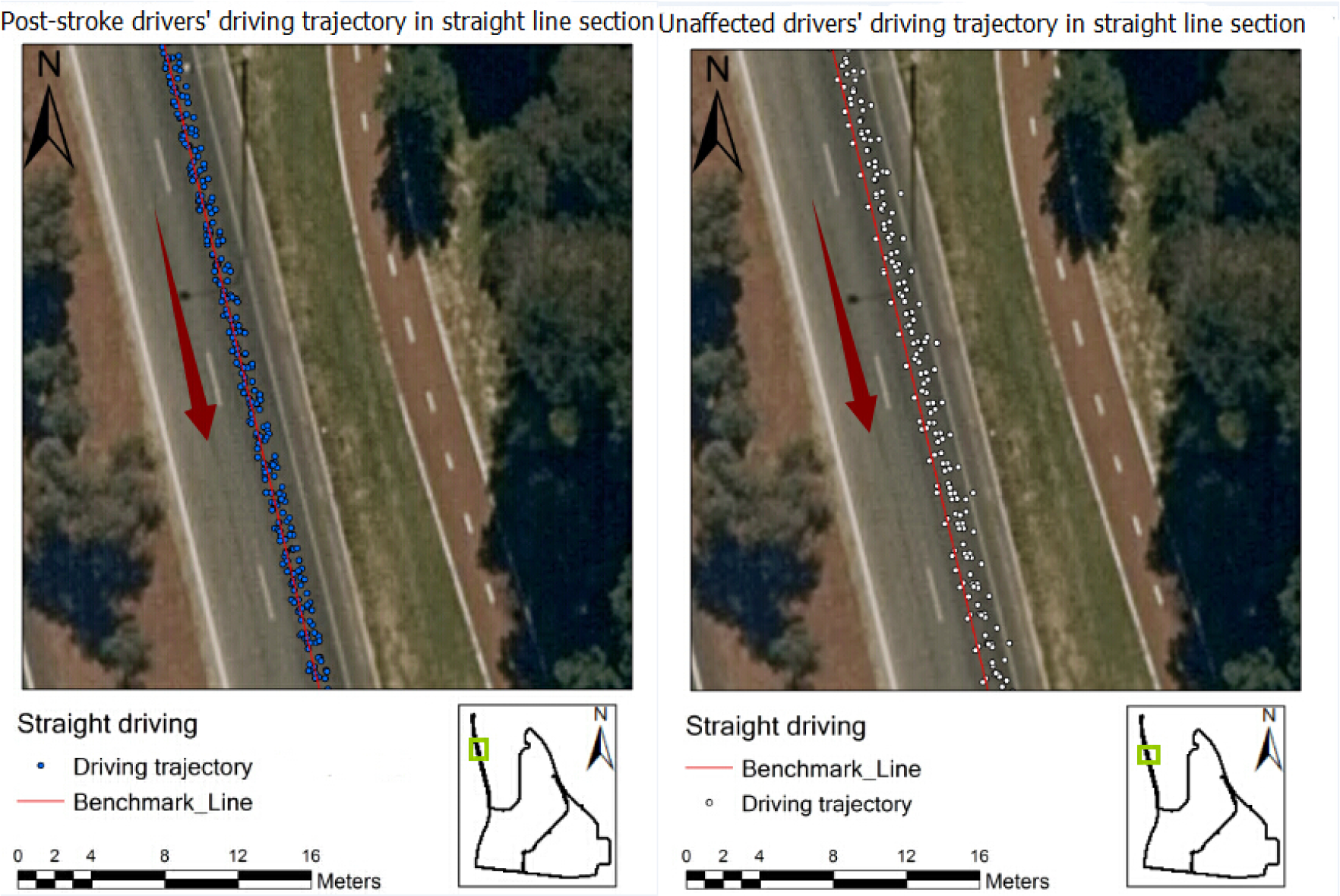
Driving trajectory of post-stroke (left) and normal (right) groups in a straight-line

**Figure 4:**
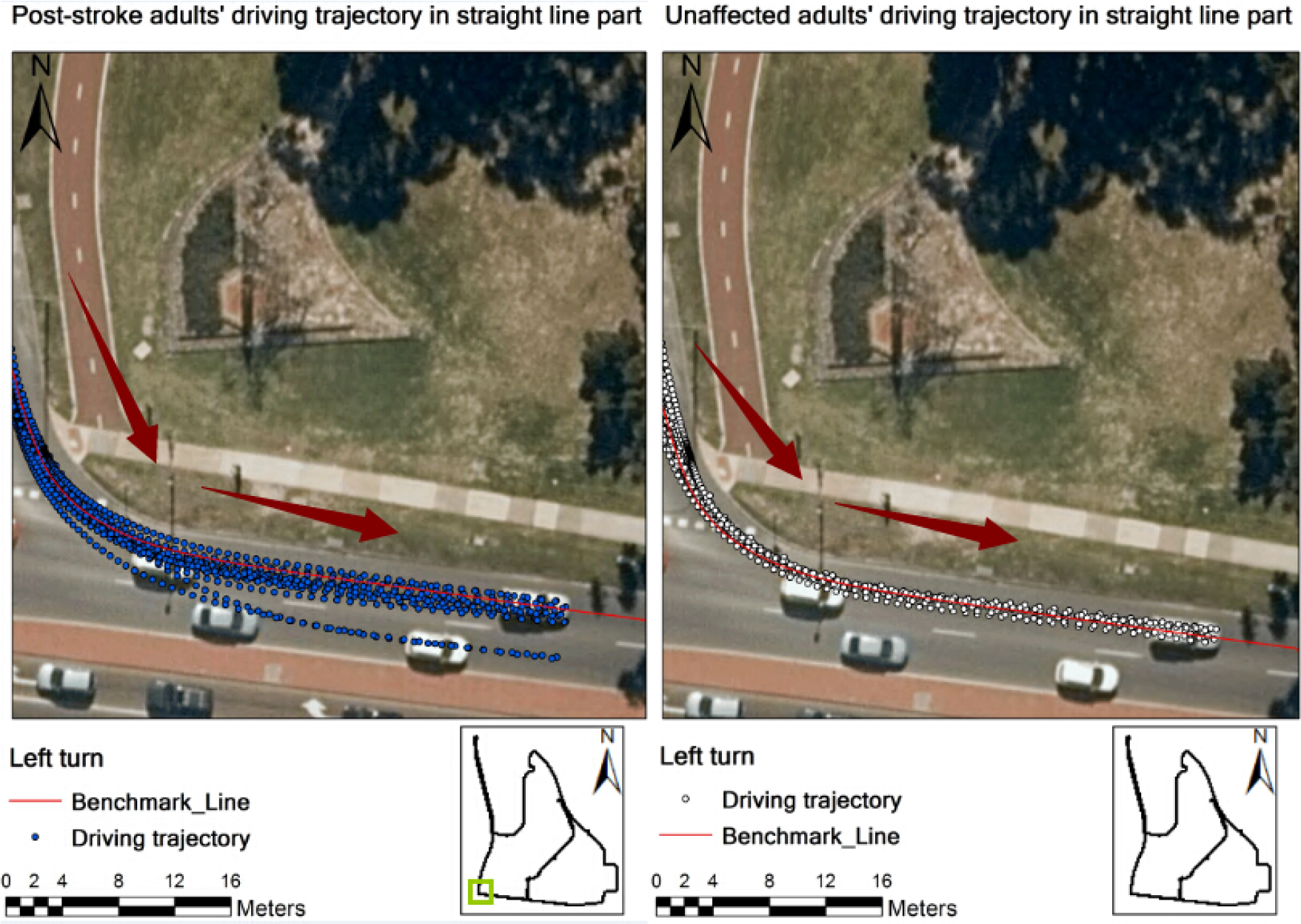
Driving trajectory of post-stroke (left) and normal (right) groups in left turn

Figure 5 shows the post-stroke drivers’ and normal drivers’ vehicle control trajectories in the Left turn of the on-road driving test. The driving trajectories exhibit that the post-stroke drivers’ deviation from road centre line in Left turn was larger than that of normal drivers. In comparison with post-stroke older drivers’ vehicle control trajectories, the normal older drivers’ vehicle control trajectories distributed more closely towards the road centre line. Thus figure 5 indicates that the post-stroke drivers’ lane maintenance in Left turns was poorer than that of normal drivers.

### 3.2 Statistical analyses of driving trajectories

#### 3.2.1 Roundabout (U-turn)

As shown in Table 1, the mean SDLD values (mean in meters) of the post-stroke group were generally higher than the values of the normal group. As outlined in table 1 in the entire U-turn, the mean SDLD of the post-stroke group was 0.60 m, which was almost twice the value of 0.32 m from the normal group; however, the *p* value was not significant. Similarly, the entry, middle part of the U-turn mean SDLD scores were generally higher, however also not significant.

In the exit section of the U-turn, the value of lane deviation in the post-stroke drivers was significantly larger than the control group’s lane deviation (1.38 m), *p*=0.043. Furthermore, in the exit of the U-turn, the mean SDLD value of post-stroke participants was 0.48 m, which was twice that of the normal group 0.19m, *p*=0.026, which suggests that the post-stroke participants’ lane keeping performance was significantly poorer than that of normal drivers.

Table 1 also shows that in U-turn exit, the post-stroke older drivers’ mean driving speed and standard deviation of speed (32.51 km/h, 5.17 km/h) are both higher than those (31.17 km/h, 4.79 km/h) of normal older drivers however differences were insignificant.

**Table 1:**
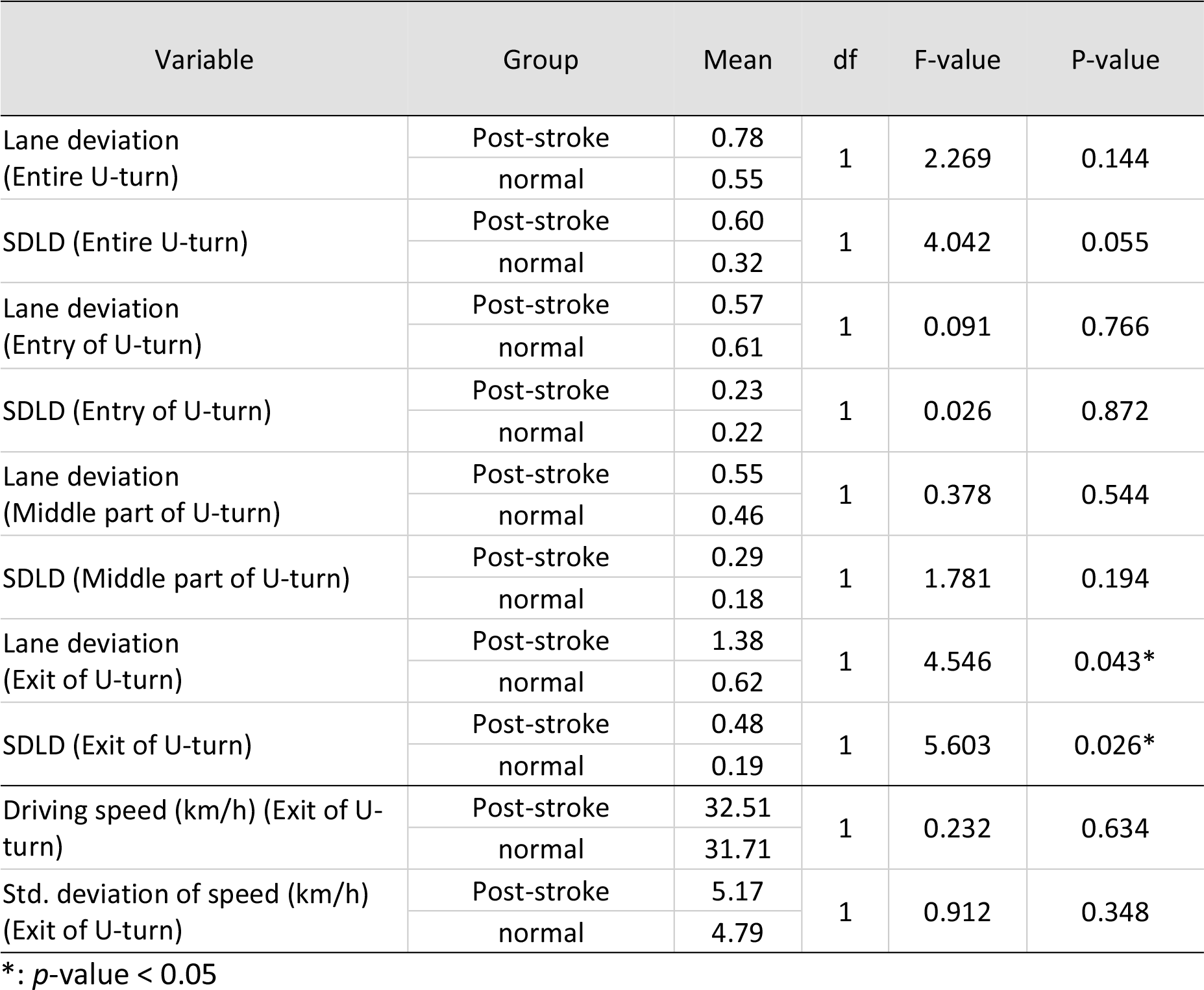
One-way ANOVA test of lane keeping performance and speed control in U-turn

#### 3.2.2 Left turn

As shown in Table 2, the mean lane deviation value (mean in meters) of the post-stroke group was 0.52 m, which was higher than that of the normal group (0.37 m). This suggests that the normal older drivers had a smaller lane deviation from the benchmark line than the post-stroke older drivers, however the differences in lane deviation were not significant. The mean SDLD of the post-stroke group was 0.38 m, which is less than that of the normal group (0.20 m) suggesting that the post-stroke older drivers may perform more poorly, but again the *p*-value (0.106) was not significant.

**Table 2:** One-way ANOVA test of lane keeping performance in left turn

| Variable | Group | Mean | df | F | P-value |
| --- | --- | --- | --- | --- | --- |
| Lane deviation (Left turn) | Post-stroke | 0.52 | 1 | 2.152 | 0.154 |
|  | normal | 0.37 |  |  |  |
| SDLD (Left turn) | Post-stroke | 0.38 | 1 | 2.796 | 0.106 |
|  | normal | 0.20 |  |  |  |

\*: $p$ -value < 0.05

#### 3.2.3 Straight line one (speed limit of 50 km/h)

As Table 3 shows, in straight line section one (speed limit of 50 km/h) the post-stroke group’s mean lane deviation value was 0.65m, which was smaller than that of the normal group (0.83 m), and the mean SDLD value of post-stroke group was 0.20m, which was also less than that of the normal group (0.28 m). Although these lower values show that the post-stroke older drivers’ maintained a more consistent line, the differences were again not significant.

The post-stroke group’s mean speed was faster than that of the normal group (48.16 and 45.00 km/h, respectively) and similarly the speed control performance measured by standard deviation of speed was greater in the post-stroke group compared to the normal group (2.58 km/h and 1.77 km/h, respectively), however the differences were not significant.

**Table 3.** One-way ANOVA test of lane keeping performance and speed control in straight line section one

| Variable | Group | Mean | df | F | P-value |
| --- | --- | --- | --- | --- | --- |
| Lane deviation (Straight line) | Post-stroke | 0.65 | 1 | 2.834 | 0.104 |
|  | normal | 0.83 |  |  |  |
| SDLD (Straight line) | Post-stroke | 0.24 | 1 | 0.987 | 0.330 |
|  | normal | 0.28 |  |  |  |
| Driving speed (km/h) | Post-stroke | 48.16 | 1 | 1.190 | 0.285 |
|  | normal | 45.00 |  |  |  |
| Std. deviation of speed (km/h) | Post-stroke | 2.58 | 1 | 2.061 | 0.163 |
|  | normal | 1.77 |  |  |  |

\*: $p$ -value < 0.05

#### 3.2.4 Straight line two (speed limit of 70 km/h)

The lane deviation and SDLD of the two groups in the straight-line section two (speed limit of 70 km/h) are summarised in Table 4. The mean lane deviation value of the post-stroke group was 0.30 m, which was half that of the normal group 0.60 m, *p*= 0.015. Furthermore, the post-stroke group’s mean SDLD value was 0.14m, which was equal to that of the normal group.

The average speed and the standard deviation of speed in post-stroke group (64.24 km/h, 1.68 km/h) was higher than that of the normal group (62.38 km/h, 1.34 km/h), but both differences were non-significant.

**Table 4:** One-way ANOVA test of lane keeping and speed control in straight line section two

| Variable | Group | Mean | df | F | P-value |
| --- | --- | --- | --- | --- | --- |
| Lane deviation (Straight line) | Post-stroke | 0.30 | 1 | 6.771 | 0.015* |
|  | normal | 0.60 |  |  |  |
| SDLD (Straight line) | Post-stroke | 0.14 | 1 | 0.049 | 0.826 |
|  | normal | 0.14 |  |  |  |
| Driving speed (km/h) | Post-stroke | 64.24 | 1 | 0.424 | 0.521 |
|  | normal | 62.38 |  |  |  |
| Std. deviation of speed (km/h) | Post-stroke | 1.68 | 1 | 1.088 | 0.307 |
|  | normal | 1.34 |  |  |  |

\*: $p$ -value < 0.05

## 4. DISCUSSION

The underlying motivation of this pilot study was to assess the feasibility, appropriateness and effectiveness of using advanced vehicle tracking (multi-GNSS RTK) and GIS analysis as a tool to differentiate driving performance of post-stroke drivers from that of normal older drivers. Subtle difference in driving competency is hard to detect using observation, but GNSS RTK technologies enable accuracy of vehicle movement tracking therefore provide detailed information on driving performance. The GNSS receiver can obtain the signal from multiple satellite systems. Additionally, RTK technology was applied after loading the trajectory data from GNSS receiver, which uses a known position of a base station to correct the recorded trajectory data and has improved the data accuracy and precision to a high level of data accuracy (sub-decimetre level). Thus, the subtle difference of the participant’s driving trajectory and behaviour was explored, which indicated that the GNSS modelling was an appropriate and effective tool to explore on-road driving performance

Although multi-GNSS RTK tracking driving behaviour has previously been utilised (Sun et al., 2017), as far as the authors are aware, this is the first study to investigate post-stroke driver performance on-road. The present study found that post-stroke adults performed more consistently in the straight-line section, and less consistently than the normal drivers in the exit to the roundabout. However, with the exception of the exit on the roundabout, there were no statistically significant differences in the speed maintenance or lane deviation performance between the post-stroke drivers and the normal adults. This would suggest that despite the differences in lane and speed deviation between groups, that the post-stroke drivers were just as safe as the control participants. This aligns well with the fact that all the post-stroke adults had been cleared to drive by a medical professional.

A small sample size is one important limitation of this study, which increases the risk of a type Ⅱ error, however as *α* value was set to 0.05, only statistical differences at the exit on the roundabout between two groups were found in the study. Further research with a greater sample size is required to address this. One may speculate that increasing the number of participants in both groups may make the difference become significant.

The gender imbalance (3 females, 11 males) in the post-stroke group is another limitation that may affect the comparison of the post-stroke and normal groups, particularly as male post-stroke are more likely to return to driving than females (McNamara, Walker, Ratcliffe, & George, 2015). However, as volunteer sampling was employed, it was difficult to recruit enough participants to gender match. Further research with a greater sample and with gender matching would address this problem.

For the on-road driving test, each participant performed the driving test using his/her own vehicle. It is possible that this may have influenced their lane keeping and driving trajectory due to tracking quality and vehicle transmission, however it can be argued that participants performance was likely to be most valid in a vehicle with which they were familiar. Also, as some participants required vehicle alterations (e.g., a spinner knob for hemiplegia or hemiparesis) for safety and insurance reasons, it was decided that all participants were to use to their own vehicle.

Although the same fixed route was applied for each participant in order to minimise the effects of road changes, one of the major difficulties in on-road testing is that the road conditions are likely change between assessments, e.g., traffic, weather, etc. In order to control for this as much as possible, participants were assessed between the hours of 10.00am – 4.00pm to avoid peak hour congestion and assessors noted whether there was any significant change in driving route, e.g., due to bad weather, road works, traffic accidents, etc.

## 5. CONCLUSIONS

It can be concluded that lane keeping might be an indicator of driving performance when driving at U turn, particularly exiting the roundabout (complex driving task), while speed control might be superior in revealing driving performance on the straight line. Cognitively demanding driving situations, such as U turn and particularly exiting the roundabout, create a challenge for the post-stroke older drivers. This may be ascribed to the higher levels of cognitive abilities required to maintain the lane along the changing geometry of the road (Cao, 2013). Understanding association between lane keeping, speed maintenance and cognitive abilities, especially divided and selective attention requires further research that might explain why some post-stroke drivers’ mean SDLD stays relatively high.

The findings of this study have demonstrated the appropriateness, feasibility and effectiveness of assessing driving behaviours of post-stroke drivers. It is strongly recommended that SDLD calculated from accurate vehicle movement trajectory is a sensitive and effective measure for driving assessment in this cohort population. Future work will need to examine and model older drivers’ lane maintenance and speed regulation in the face of hazardous driving situations. Further educational and training programs based on the findings of this study could be developed to enhance post-stroke drivers’ behaviour behind the wheel; for example, neuropsychological training to improve the post-stroke drivers’ executive function (Rand & Katz, 2009); and driving intervention training to improve lane keeping performance (Sawula et al., 2018) at challenging driving sections.

## APPENDIX A

The on-road driving assessment route with focus junctions and base stations indicated

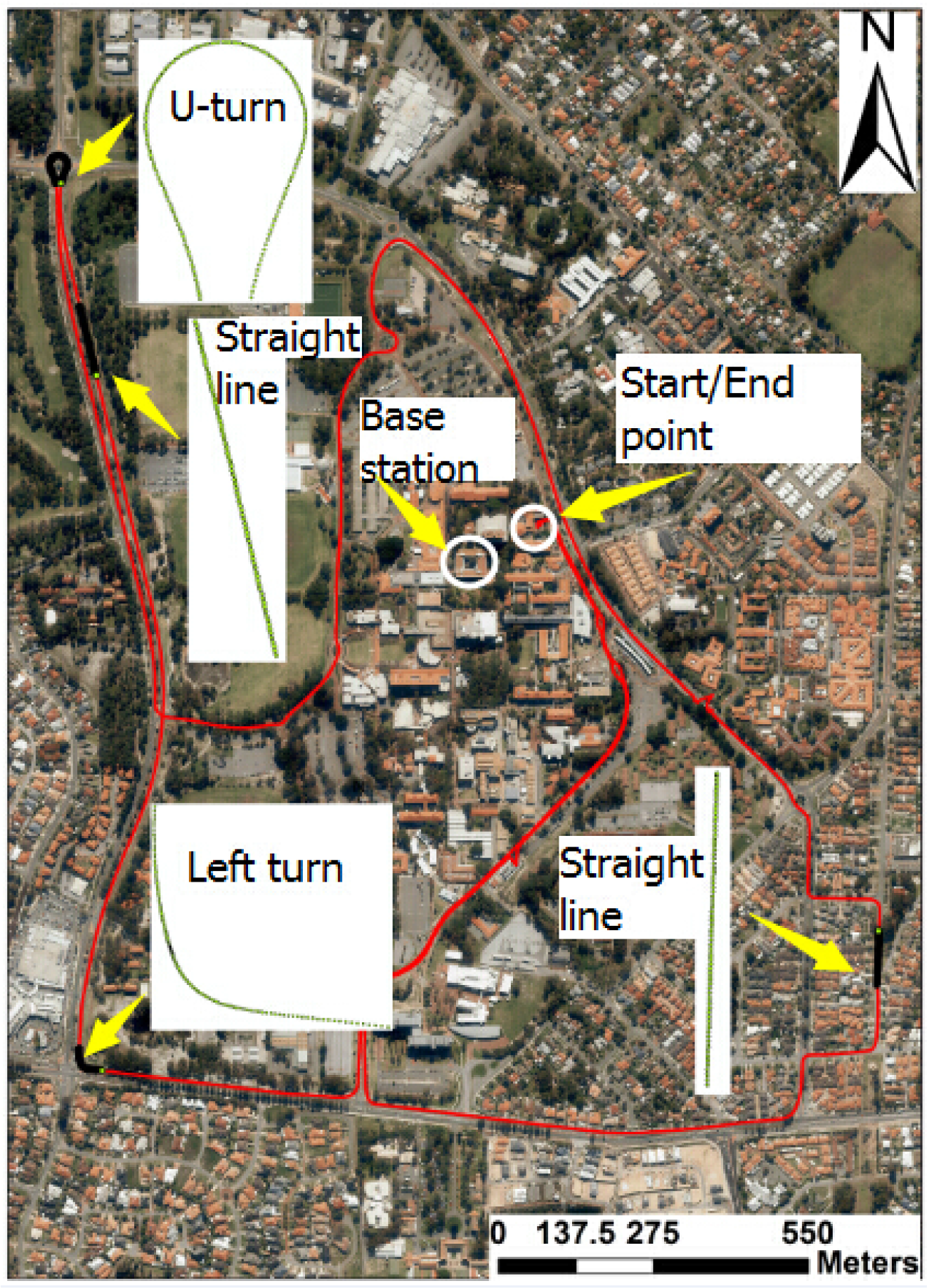

## REFERENCES

Anstey, K. J., Wood, J., Lord, S., & Walker, J. G. (2005). Cognitive, sensory and physical factors enabling driving safety in older adults. Clinical Psychology Review.

Burns, P. C. (1999). Navigation and the mobility of older drivers. J. Gerontol. Ser. B-Psychol. Sci. Soc. Sci., 54(1), S49–S55.

Cao, S., & Liu, Y. (2013a). Concurrent processing of vehicle lane keeping and speech comprehension tasks. Accident Analysis and Prevention, 59, 46–54. doi:10.1016/j.aap.2013.04.038

Cao, S., & Liu, Y. (2013b). Concurrent processing of vehicle lane keeping and speech comprehension tasks. Accident Analysis & Prevention, 59, 46–54. doi:http://dx.doi.org/10.1016/j.aap.2013.04.038

Carsten, O., Kircher, K., & Jamson, S. (2013). Vehicle-based studies of driving in the real world: The hard truth? Accident Analysis and Prevention, 58, 162–174. doi:10.1016/j.aap.2013.06.006

Chan, E., Pradhan, A. K., Pollatsek, A., Knodler, M. A., & Fisher, D. L. (2010). Are driving simulators effective tools for evaluating novice drivers’ hazard anticipation, speed management, and attention maintenance skills? Transportation Research Part F: Traffic Psychology and Behaviour, 13(5), 343–353. doi:http://dx.doi.org/10.1016/j.trf.2010.04.001

El-Rabbany, A. (2006). Introduction to GPS – The Satellite Navigation System (Second ed.). Artech House, Norwood, MA

Harbluk, J. L., Noy, Y. I., & Eizenman, M. (2002). The impact of cognitive distraction on driver visual behaviour and vehicle control (00942271). Retrieved from Ontario, Canada: http://www.tc.gc.ca/roads…/tp13889/pdf/tp13889es.pdf

He, J., McCarley, J. S., & Kramer, A. F. (2014). Lane Keeping Under Cognitive Load. Human Factors: The Journal of Human Factors and Ergonomics Society, 56(2), 414–426. doi:10.1177/0018720813485978

Howard, G., & Goff, D. C. (2012). Population shifts and the future of stroke: forecasts of the future burden of stroke. Ann.NY Acad.Sci., 1268, 14–20. doi:10.1111/j.1749-6632.2012.06665.x

IBM Corporation. (2012). IBM SPSS Statistics for Windows, Version 21.0. Armonk, NY: IBM Corp.

Mathias, J. L., & Lucas, L. K. (2009). Cognitive predictors of unsafe driving in older drivers: A meta-analysis. International Psychogeriatrics, 21(04), 637–653.

McGwin, G., Sims, R. V., Pulley, L., & Roseman, J. M. (2000). Relations among Chronic Medical Conditions, Medications, and Automobile Crashes in the Elderly: A Population-based Case-Control Study. American Journal of Epidemiology, 152(5), 424–431. doi:10.1093/aje/152.5.424

McNamara, A., Walker, R., Ratcliffe, J., & George, S. (2015). Perceived confidence relates to driving habits post-stroke. Disability & Rehabilitation, 2015, Vol.37(14), p.1228–1233, 37(14), 1228-1233. doi:10.3109/09638288.2014.958619

Motta, K., Lee, H., & Falkmer, T. (2014). Post-stroke driving: Examining the effect of executive dysfunction. Journal of Safety Research, 49, 33.e31. doi:10.1016/j.jsr.2014.02.005

National stroke foundation. (2014). Supports limiting life after stroke New report. Retrieved from https://strokefoundation.com.au/news/2015/05/18/supports-limiting-life-after-stroke-new-report

Perrier, M.-J., Korner-Bitensky, N., Petzold, A., & Mayo, N. (2010). The risk of motor vehicle crashes and traffic citations post stroke: A structured review. Topics in Stroke Rehabilitation, 17(3), 191–196.

Poole, D., Chaudry, F., & Jay, W. M. (2008). Stroke and driving. Topics in Stroke Rehabilitation, 15(1), 37–41.

Rabadi, M., Akinwuntan, A., & Gorelick, P. (2010). The Safety of Driving a Commercial Motor Vehicle After a Stroke. Stroke, 41(12), 2991–2996.

Rabadi, M. H., Akinwuntan, A., & Gorelick, P. (2010). The Safety of Driving a Commercial Motor Vehicle After a Stroke. Stroke, 41(12), 2991–2996. doi:10.1161/strokeaha.110.587196

Rakotonirainy, A., Steinhardt, D., Delhomme, P., Darvell, M., & Schramm, A. (2012). Older drivers’ crashes in Queensland, Australia. Accident Analysis & Prevention, 48(0), 423–429. doi:http://dx.doi.org/10.1016/j.aap.2012.02.016

Reed, M. P., & Green, P. A. (1999). Comparison of driving performance on-road and in a low-cost simulator using a concurrent telephone dialling task. Ergonomics, 42(8), 1015–1037. doi:10.1080/001401399185117

Selander, H., Falkmer, T., & Bolin, I. (2012). Does automatic transmission improve driving behavior in older drivers? Gerontology, 58(2), 181–187. doi:10.1159/000329769

Selander, H., Lee, H., Johansson, K., & Falkmer, T. (2011). Older drivers: On-road and off-road test results.

Sun, Q., Xia, J., Nadarajah, N., Falkmer, T., Foster, J., & Lee, H. (2016). Assessing drivers’ visual-motor coordination using eye tracking, GNSS and GIS: a spatial turn in driving psychology. Journal of Spatial Science, 1–18.

Sun, Q., Xia, J. C., Foster, J., Falkmer, T., & Lee, H. (2016). Pursuing Precise Vehicle Movement Trajectory in Urban Residential Area Using Multi-GNSS RTK Tracking. Paper presented at the World Conference on Transport Research – WCTR 2016, Shanghai, China.

Sun, Q. C., Odolinski, R., Xia, J. C., Foster, J., Falkmer, T., & Lee, H. (2017). Validating the efficacy of GPS tracking vehicle movement for driving behaviour assessment. Travel Behaviour and Society, 6, 32–43.

Sun, Q. C., Xia, J. C., Foster, J., Falkmer, T., & Lee, H. (2018a). Driving manoeuvre during lane maintenance in older adults: Associations with neuropsychological scores. Transportation Research Part F: Traffic Psychology and Behaviour, 53, 117–129.

Sun, Q. C., Xia, J. C., Foster, J., Falkmer, T., & Lee, H. (2018b). A psycho-Geoinformatics approach for investigating older adults’ driving behaviours and underlying cognitive mechanisms. European Transport Research Review, 10(2), 36.

Terblanche, W., & Wilson, T. (2015). An Evaluation of Nearly-Extinct Cohort Methods for Estimating the Very Elderly Populations of Australia and New Zealand. PLoS One, 10(4). doi:10.1371/journal.pone.0123692

Weihong, G., Blythe, P. T., Edwards, S., Pavkova, K., & Brennan, D. (2015). Effect of intelligent speed adaptation technology on older drivers’ driving performance. Intelligent Transport Systems, IET, 9(3), 343–350. doi:10.1049/iet-its.2013.0136

White, R. W. (1959). Motivation reconsidered: The concept of competence. Psychological Review, 66(5), 297–333. doi:10.1037/h0040934

